# Proteomic Dissection of a Giant Cell

**DOI:** 10.1101/2022.01.06.475287

**Authors:** Athena Lin, Paul D. Piehowski, Chia-Feng Tsai, Tatyana Makushok, Lian Yi, Ulises Diaz, Connie Yan, Diana Summers, Pranidhi Sood, Richard D. Smith, Tao Liu, Wallace F. Marshall

## Abstract

Many individual proteins have been identified as having defined positions relative to cell polarity axes, raising the question of what fraction of all proteins may have polarized localizations. We took advantage of the giant ciliate *Stentor coeruleus* to quantify the extent of polarized localization proteome-wide. This trumpet-shaped unicellular organism shows a clear morphological anterior-posterior axis defined by a circular array of cilia known as a membranellar band at one end, and a holdfast at the other end. Because individual *Stentor* cells are over a millimeter in length, we were able to cut the cells into three pieces along the anterior-posterior axis, followed by proteomic analysis of proteins enriched in each piece. We find that approximately 30% of all detected proteins show a polarized location relative to the anterior-posterior cell axis. Proteins with polarized enrichment include centrin-like proteins, calcium-regulated kinases, orthologs of SFI1 and GAS2, and proteases. At the organelle level, nuclear and mitochondrial proteins are enriched in the anterior half of the cell body, but not in the membranellar band itself, while ribosome related proteins are apparently uniformly distributed. RNAi of signaling proteins enriched in the membranellar band, which is the anterior-most structure in the cell, revealed a protein phosphatase 2 subunit b ortholog required for closure of the membranellar band into the ring shape characteristic of *Stentor.* These results suggest that a large fraction of the *Stentor* proteome has a polarized localization, and provide a protein-level framework for future analysis of pattern formation and regeneration in *Stentor* as well as defining a general strategy for subcellular spatial proteomics based on physical dissection of cells.

## Introduction

Cells are not simply amorphous bags of enzymes, but rather are highly structured molecular systems, in which different molecules are partitioned into compartments which, themselves, are non-randomly arranged with respect to the shape and polarity of the whole cell (Kirschner and Mitchison 2000; Shulman and St. Johnston 2000; Harold 2005; Marshall 2011). At the level of organelles and multi-molecular structures, cells often show a startling degree of patterning, including anterior-posterior and apical-basal polarization axes and, in some case, left-right asymmetry (Frankel 1989). Patterning within cells extends down to the level of proteins and mRNA (Shulman and St. Johnston 1999; Das 2021). During the development of many organisms, non-random localization of cell-fate determinants in the egg plays a key role in dictating the pattern of subsequent development in the cleavage products of the fertilized egg (Grunert 1996). Such developmentally-relevant segregation of fate determinants is also a common feature of stem cell differentiation (Knoblich 2010, Venkei 2018). Nonrandom protein localization within cells plays a key role in basic cell biological phenomena such as motility, secretion, cell division, and cell-cell interaction. Understanding the distribution of proteins within a cell is thus a central issue for both cell and developmental biology. Perhaps the most fundamental question is simply: how much of the proteome is localized? Do all or most proteins have a spatially restricted localization relative to cell polarity axes, or are polarized proteins just a small subset of the total proteome?

Proteomics has been combined with cellular fractionation to map proteins onto organelles (Foster 2006; Dunkley 2006; Itzhak 2016; Pandya 2017). These studies have found that a large fraction of proteins localize to one or more organelle, but because spatial information is lost during the fractionation, they do not directly address the question of how many proteins have a polarized localization with respect to the large-scale polarity axes of the intact cell. Systematic immunofluorescence (Thul 2017) in mammalian cells obtained a similar result to that seen with proteomics, but did not address the polarization question because the cell types reported do not have a clear polarity axis.

Genome-wide localization studies in yeast using epitope or GFP tagging (Kumar 2002; Huh 2003; Narayanaswamy 2009; Chong 2015;) have found that the majority of proteins (70%) show clear localization to one or more organelles or other cellular structures, while approximately 30% of proteins show a more uniform distribution in the cytoplasm. In these studies, however, only a few percent of proteins appear to have a localized position relative to the cell polarity axis defined by sites of polarized cell growth, raising the possibility that only a similarly small fraction of proteins may be polarized in other cell types.

Here we describe an approach for analyzing protein distribution within a cell, based on a combination of physical dissection and proteomics in the giant ciliate *Stentor coeruleus. Stentor* is a classical model system for the study of cell morphogenesis and regeneration (Tartar 1961; Marshall 2021). Ciliates, including *Stentor,* have highly polarized cell shapes with easily visualized surface patterning, making them ideal systems in which to map proteins relative to clearly visible axes of polarity (Aufderheide 1980; Frankel 1989). *Stentor* (**Figure 1**) is particularly advantageous because of its huge size - a single cell can grow up to 2 mm in length, comparable in size to many embryos or small animals. The huge size of a *Stentor* cell means that even a small piece of the cell will contain sufficient protein for detection by mass spectrometry methods.

**Figure 1.**
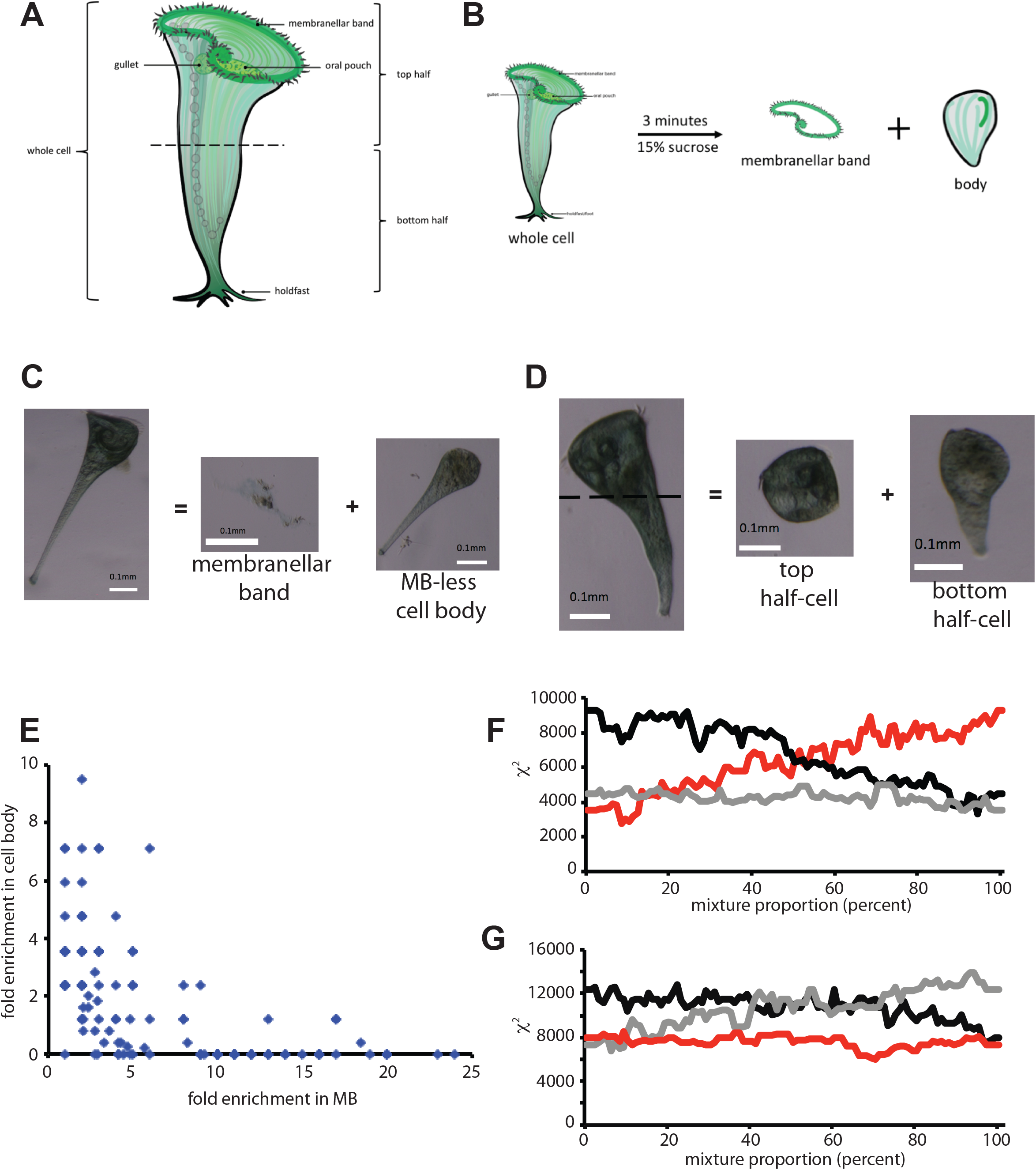
Proteomic dissection of *Stentor.* (A) Diagram of *Stentor* cell showing major anatomical features. (B) Removal of membranellar band (MB) by sucrose shock. (C) Example of a cell before and after sucrose shock, showing detachment of membranellar band and the residual MB-less cell body. (D) Manual dissection of a *Stentor* cell by cutting with a glass needle to produce anterior and posterior half cells. (E) correlation between enrichment in MB and depletion from body. Enrichment is reported as fold enrichment in each fraction relative to intact cells, such that a fold enrichment less than 1.0 indicates net depletion in the cell fraction. (F) Dissection Profile analysis of MB removal as described in Methods. (red) mixture model p*bodies + (1-p)*MB = whole cells, plotting chi squared for sum compared to whole cells, as a function of percent p of MB. (black) control mixture model pMB+(1-p)whole cells = bodies. (grey) control mixture model pWhole cells + (1-p)bodies = MB. (G) Dissection Profile analysis of cell bisection. (red) mixture model p*anterior + (1-p)*posterior = whole cells. (black) control mixture model p*whole cells + (1-p)*anterior = posterior. (grey) control mixture model p*posterior + (1-p)*whole cells = anterior. The identities of proteins detected in our analysis are tabulated in **Table S1**.

By performing “proteomic dissection” of *Stentor,* using microsurgery to separate cellular fragments along the anterior-posterior axis followed by comparative proteomic analysis, we find that approximately 30% of the proteome shows a polarized location along the cell’s anterior-posterior axis. These localized proteins, which include signaling proteins, centrin/SFI proteins, and multiple GAS2 orthologs, provide a molecular entry point to future analysis of pattern formation and regeneration within a single cell.

## Results

### Comparative proteomic analysis of the anterior-posterior axis of *Stentor*

The anterior-posterior axis of the *Stentor* cell is defined by a membranellar band (MB) consisting of a dense array of motile cilia at one end of the cell, which is defined to be the anterior end, and a holdfast at the other end, which is defined as posterior. Between these two structures, other cellular structures such as the macronucleus or contractile vacuole typically show defined positions. We thus chose the anterior-posterior axis as the reference for testing our proteomic dissection strategy, because of these visible markers. We physically dissected cells into three sectors (**Figure 1A**): membranellar band, anterior half body, and posterior half body. The membranellar band is easily dissociated from the cell using a sucrose shock (Lin 2018). The cell body itself was cut in half using a glass needle. Each sample was hand-isolated into different collection tubes. In total, five samples of Stentor were analyzed using mass spectrometry: whole cells, bodies (cells that have shed their MB), MB, top halves and bottom halves. A total of 1754 proteins were detected at least twice among the 5 samples (**Table S1)**.

We first consider the MB dataset to ask if it contains expected MB proteins, while lacking proteins we expect to not be present. The most abundant proteins identified in our proteomic analysis of the MB fraction (**Table S1**) were tubulin and axonemal dynein. We confirmed this result biochemically. On a polyacrylamide gel (**Supplemental Figure S1A**) isolated MB showed two dominant bands which we cut and analyzed by mass spectrometry. The band at 55 kDa was found to contain both alpha and beta tubulin (SteCoe_14841 and SteCoe_22851), while the larger band at 250 kDa primarily contained axonemal inner dynein arm heavy chain (SteCoe_11197). The tubulin band was further confirmed by Western blotting (**Supplemental Figure S1B**).

When MB are shed by sucrose shock, the macronucleus is retained in the cell body. Consequently, histone proteins, which are abundant in the cell, should not be abundant in the MB. Our proteomic analysis detected 12 histone proteins (**Table S1**). 7 of these proteins were not detected at all in the MB sample, and the remainder were present at substantially lower quantities than in the other cellular fragments. Based on median-normalized counts, histones were enriched approximately 5-fold in cell bodies lacking MB compared to the isolated MB. Similar results were obtained in analyses of mitochondrial and ribosomal contamination. Out of 131 proteins annotated as involved in translation (mostly ribosomal proteins or translation elongation factors), that were detected in either the MB or the headless bodies, the relative enrichment (relative to whole cells) was greater for headless bodies than for MB in 113 of the 131 proteins. Out of 40 annotated mitochondrial proteins, 35 were detected in either the MB or the body dataset. Out of those 35, the vast majority (29 proteins) were more enriched in the headless bodies relative to whole cells compared to the MB. 22 of the 40 mitochondrial proteins were not detected in the MB at all. Out of the 10 most abundant mitochondrial proteins, 9 were more enriched in the headless bodies than in the MB fraction.

Taken together these observations indicate that expected MB proteins were detected as abundant in the MB fraction, while known non-MB proteins were reported as enriched in headless bodies compared to the MB fraction, thereby providing confirmation that our proteomic results matched prior expectations.

Transcriptomic data provide a second way to evaluate specificity of proteomic composition of the MB. We previously reported an RNA sequencing study of gene expression during MB regeneration, and it is to be expected that many of the upregulated genes should encode protein components of the MB. **Table S1** includes the results of our prior RNAseq study for proteins detected in our proteomic analysis. Out of the 1754 proteins reported in our combined proteome of all samples, 281 corresponded to genes upregulated during regeneration. Among these, the average median centered abundance was 4.4 +/− 0.5 SEM for the MB sample, compared to 0.9 +/− 0.1 SEM for cell bodies lacking MB. For comparison, among the remaining 1473 proteins that did not correspond to genes upregulated during OA regeneration, the average abundance was 1.6 +/− 0.2 and 2.4 +/− 0.1, respectively. Thus, proteins whose genes are upregulated during MB regeneration are enriched in our MB proteome dataset. Conversely, of 433 proteins that are enriched at least ten-fold in the MB compared to MB-less bodies, 164 correspond to genes upregulated in cells regenerating the MB (38%). Out of the remaining 1322 proteins, only 118 are upregulated (9%). This correlation between differential proteomic enrichment in the MB and upregulation during MB regeneration, supports the conclusion that our analysis is detecting bona fide MB-related proteins.

In contrast to the MB, we have less prior knowledge of proteins expected to be present in the anterior versus posterior half cells. It is, however, known that the posterior of the cell contains extensive contractile fibers composed of centrin-related EF hand proteins (Maloney 2005). Consistent with this expectation, of the five centrin proteins identified in the proteome, all five were more abundant in the posterior half compared to the anterior half or to the MB (**Table S1**).

The anterior half-cell contains the MB, and should thus show an enrichment for MB-specific proteins compared to the posterior half. By comparing the relative enrichment of proteins in the MB fraction vs whole cells with the relative enrichment of proteins in the anterior half-cell vs whole cells, limiting the analysis to abundant proteins (at least 10 hits), we found a significant correlation (r=0.33; (p<0.0001, n=481) confirming the anterior fraction includes MB proteins.

As a final test of selectivity in the anterior vs posterior half-cell experiment, we compared differential gene expression in the anterior and posterior halves of bisected Stentor cells (Sood 2021) to our differential protein abundance data. When cells are bisected, posterior half-cells regenerate new anterior structures such as the MB, while anterior half cells regenerate new posterior structures such as the holdfast. Of three genes upregulated during regeneration of new posterior structures by anterior half-cells, that also corresponded with proteins detected in our proteomic analysis, the average relative abundance in the posterior half versus the anterior half was 23-fold, compared to an average of 1.4 fold taken over all proteins. This is a small number of genes but it confirms that the posterior cell fraction is likely enriched for posterior localized proteins.

As a further way to validate our results, we considered two global criteria for successful proteomic dissection. First, proteins enriched in any given cell sector are expected to be depleted from the remainder of the cell that is missing that sector. If, instead, differences between samples were due to random variation in protein detection, samples should be uncorrelated with each other. We selected all proteins enriched at least two-fold in the MB versus whole cells, and compared the enrichment in the MB vs whole cell to the relative enrichment in cell bodies from which the MB had been removed vs whole cell (**Figure 1E**). We observed a significant negative correlation (r=-0.34; n=119; P<0.0002) confirming that proteins enriched in one fraction are depleted from the complementary fraction.

As a second global criterion for successful proteomic dissection, we used a mixture model to ask whether the combined protein abundance profiles from complementary pairs of cell fractions could recapitulate protein abundance profiles consistent with those seen in the whole cell. The results of this analysis, described in detail in Materials and Methods, is shown in **Figure 1F and G**, which show that for both the MB vs bodies, and for the cell bisection comparison of anterior versus posterior halves, a combination of both fragments match the protein profile of whole cells better than either fragment alone.

### Quantifying the fraction of proteins with polarized localization

A major question for subcellular proteomics is to what extent are proteins non-uniformly distributed within a cell. **Figure 2A** plots the correlation coefficient between the relative abundances (normalized by whole cell data), between membranellar bands, cell bodies missing the MB, anterior halves of bisected cells, and posterior halves of bisected cells. This plot indicates that the MB fraction shows a low correlation with all the other fractions (blue bars), supporting the idea that the proteome of the MB is distinct from the rest of the cell. Consistent with results given above, the correlation coefficient between MB and cell bodies lacking the MB is negative. The cell body fraction is correlated with both the anterior and posterior fractions (red bars), while the highest correlation was seen between the anterior fraction and the posterior fraction (green bar). This latter result suggests that a large number of proteins in the cell body are equally distributed among the anterior and posterior fractions. This raises the question - besides the MB enriched proteins, how many proteins, if any, show enrichment in the anterior versus posterior cell body?

**Figure 2.**
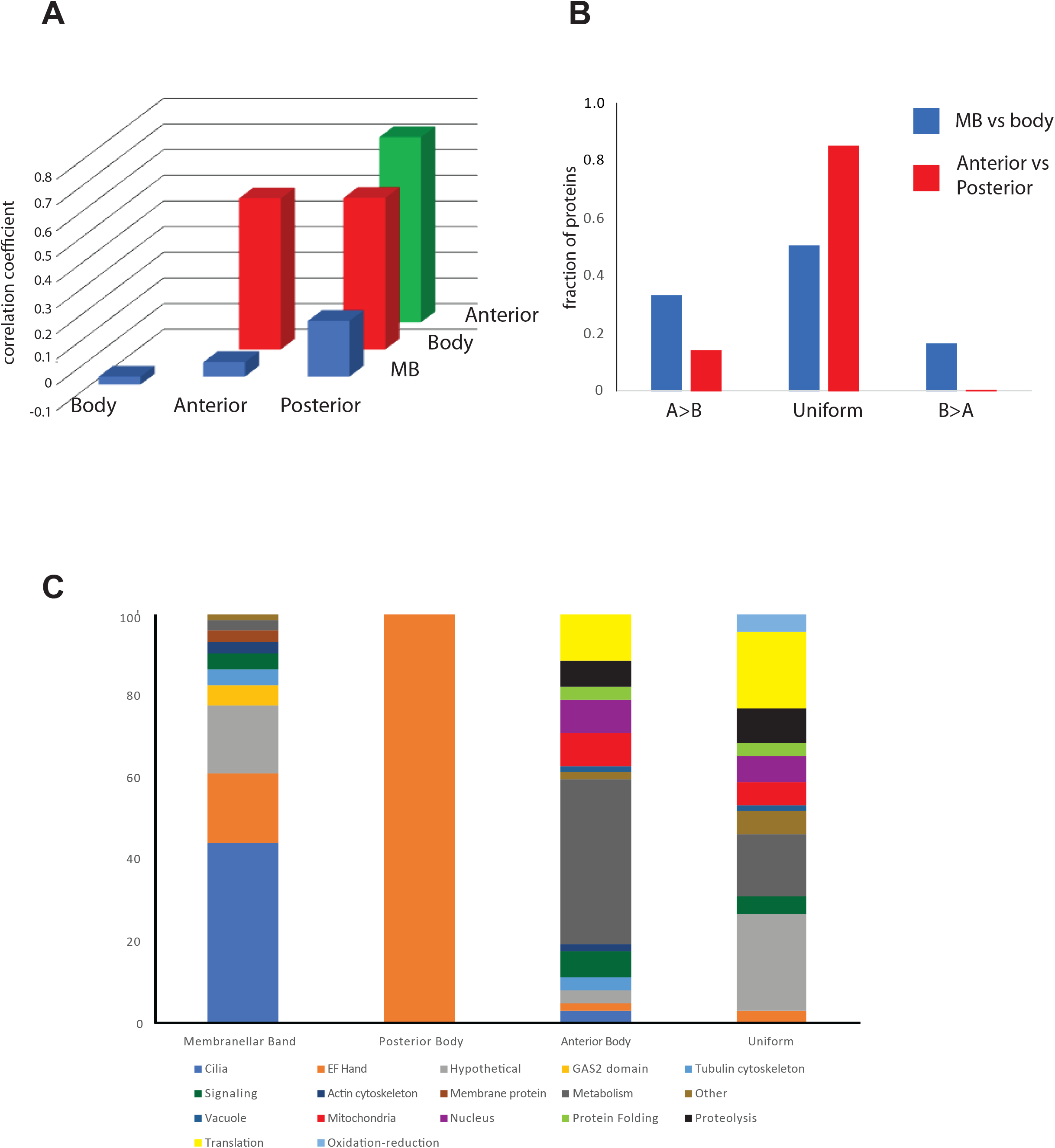
Global analysis of protein distribution. (**A**) Analysis of correlation between enrichment in different cell fractions. Bars indicate the correlation coefficient in relative abundance (median normalized abundance in the given fraction relative to whole cells) between each of two fractions as indicated on the XY axes. Body refers to cell bodies from which the MB has been removed. (**B**) Proportion of proteins significantly enriched (p<0.05) in one compartment versus another based on binomial test. The category A>B indicates proteins that are significantly more abundant in the first versus second sample as indicated in the legend (blue: more abundant in MB than in cell body red: more abundant in anterior than in posterior). Uniform indicates proportion of proteins showing no significant different in abundance between the two samples being compared. B>A indicates proteins that are significantly more abundant in the second sample (cell body or posterior) than in the first sample (MB or anterior) for each of the two comparisons. (**C**) Protein families represented in the differentially enriched gene sets described in panel B.

In order to evaluate the level of protein non-uniformity, we assessed the degree to which proteins are non-uniformly localized by comparing two pairs of samples. First, we compared the MB fraction to cell bodies from which the MB had been removed. We used a binomial test to identify proteins that were significantly enriched in one fraction versus the other (see Methods) at a significance value of 0.05. Out of 398 proteins with at least five total hits between the MB and body, 65 are enriched in the body, 133 are enriched in the MB, and 200 are uniformly distributed between the two fractions (**Figure 2B**, blue bars). We next considered non-MB enriched proteins and asked whether they showed nonuniform localization between the anterior and posterior half-cells as described in Methods. Out of 443 non-MB enriched proteins with at least five hits between anterior and posterior, 63 are significantly enriched in the anterior relative to posterior, 3 are enriched in posterior relative to anterior, and the remaining 377 are uniformly distributed between the two fractions (**Figure 2B**, red bars).

The plot in **Figure 2B** compares localization in MB versus body separately from localization in anterior versus posterior bodies. If we consider all proteins detected with at least 5 hits among all fractions (916 proteins total- see **Table S1**), we find that of 916 proteins, 3 are enriched in the cell body posterior, 97 are enriched in the cell body anterior but not in the MB, 103 are enriched in the MB but not in the cell body anterior, and 122 are enriched in both the cell body anterior and the MB, for a total of 325 proteins enriched in either the MB, cell body anterior, or cell body posterior, which constitutes 35% of the total proteins analyzed. Given that our criterion for enrichment is defined by a binomial test at the 5% significance level, we conclude that approximately 30% of the proteome shows polarized localization.

**Figure 2C** indicates the protein families contained in the differentially enriched groups from **Figure 2B**. The MB enriched protein set clearly contains many cilia proteins as well as GAS2 domain proteins. The posterior body enriched protein set is dominated by EF hand related proteins, in particular orthologs of centrin and the centrin-organizing protein SFI1. The anterior body enriched protein set contains a large number of mitochondrial and nuclear proteins. While SFI1 family proteins, known to be involved in organizing centrin filaments in other organisms, are found in both the anterior and posterior of the cell, they are different specific proteins, such that there are distinct MB/anterior specific and posterior-specific SFI1 proteins. In addition to these polarized proteins, other EF-hand proteins were found to be uniformly distributed. The results of this analysis are consistent with a simpler pairwise analysis of proteins enriched in individual fractions relative to other individual fractions, as shown in **Supplemental Figure S2**.

### Identifying polarized proteins in the cell body

In our analysis of anterior versus posterior half cells, the presence of the membranellar band was a potentially confounding factor in searching for proteins localized in the anterior half of the cell body. Although we attempted to avoid this effect in **Figure 2B** by removing MB enriched proteins computationally from the analysis of anterior vs posterior halves, the influence of the MB on the comparison remained a potential concern. Furthermore, both anterior and posterior half-cells contained a shared set of highly abundant proteins such as tubulin, which tended to account for a large fraction of all spectral counts, making it difficult to detect less abundant proteins that might show interesting regional differences. We therefore searched for less abundant proteins that might be enriched in one half of the cell versus the other, by bisecting cell bodies after removal of the MB (**Figure 3A**) and then performing mass spectrometry using tandem mass tagging (TMT) to label proteins from the two fractions allowing them to be analyzed side by side (**Figure 3B**). This analysis revealed 97 proteins that were enriched in the anterior half body compared to the posterior, and 109 proteins enriched in the posterior compared to the anterior (**Figure 3C**). The results of the analysis described in **Figure 3** are given in **Table S2**. Among the proteins enriched in the anterior and posterior fractions according to this analysis were GAS2 and centrin/SFI related proteins respectively. 12/12 mitochondria proteins detected in the TMT analysis were enriched in the anterior half. These results are therefore consistent with the comparative proteomic data of **Figure 2C**.

**Figure 3.**
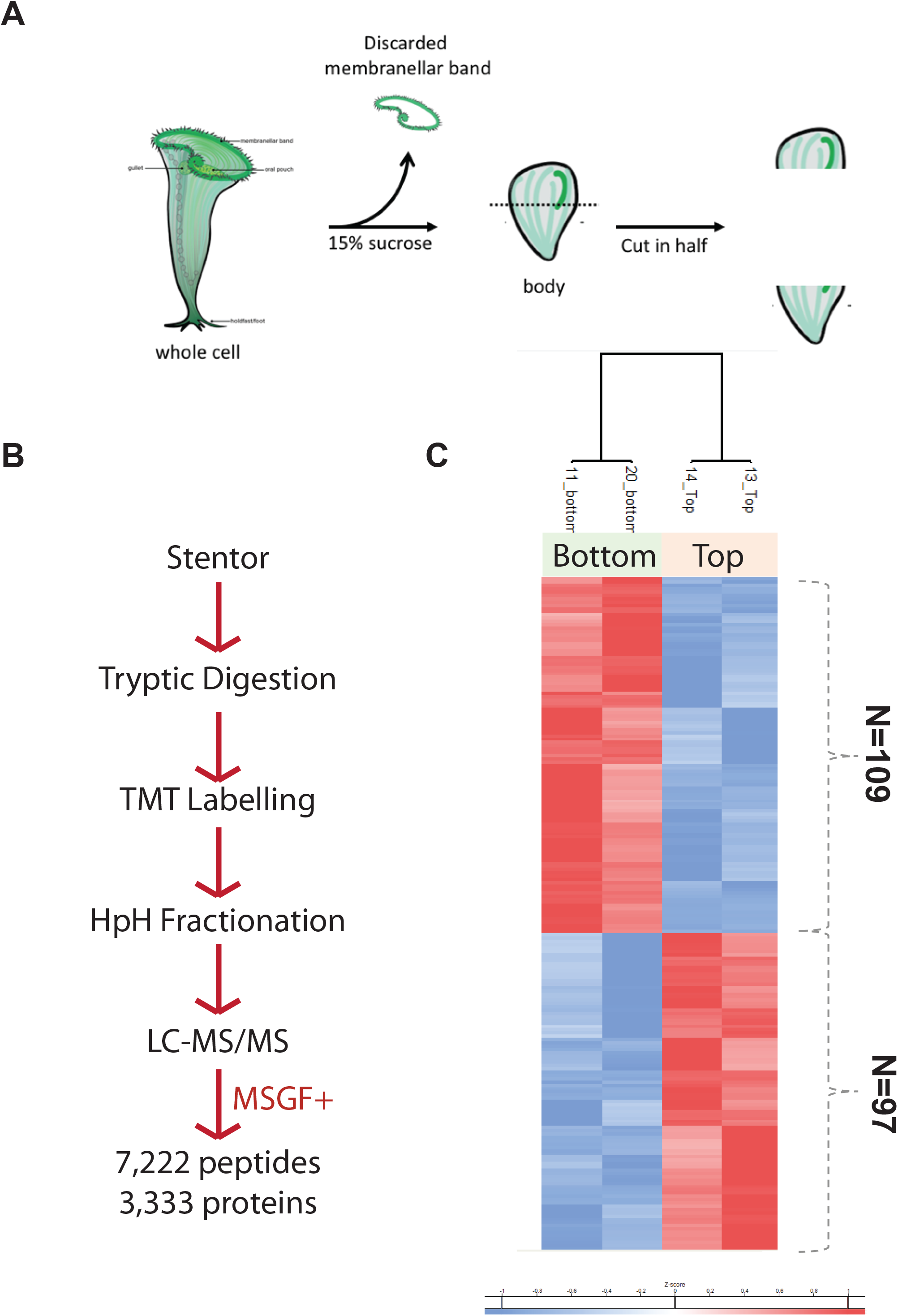
Proteomic comparison between anterior and posterior cell bodies. (**A**) In order to prevent MB proteins from dominating the protein comparison, the MB was first removed by sucrose shock, after which the MB-less cell body was cut in half. (**B**) Proteomic flow chart for differential comparison of cell body halves. TMT refers to tandem mass tagging the peptides by reacting them with isobaric tags. MSGF+ refers to computational search and identification of isobaric tagged peptides using the MS-GF+ software. (**C**) Cluster diagram showing proteins enriched in the anterior and posterior halves of the cell body. Proteins depicted in this heatmap are listed in **Table S2**.

### Proteome of soluble fraction from membranellar bands

In considering proteins localized to the membranellar band, we were particularly interested in soluble proteins, in light of previous suggestions that MB regeneration is triggered by loss of a diffusible signal generated within the MB when it is present (Hyvert 1972). The samples analyzed above were solubilized for mass spectrometry using a chemical procedure designed to maximally disrupt proteinprotein interactions, chosen because the ciliate cortex is known to consist of a highly stable protein network. In order to specifically analyze soluble MB proteins, hand-isolated MB, as well as intact cell bodies and cell bodies after MB removal, were treated with a gentle detergent lysis procedure, the insoluble material pelleted, and the supernatant retained for analysis by mass spectrometry. Proteins detected in these three samples are listed in **Table S3**. The correlation coefficient for enrichment in MB and depletion from MB-less bodies among the top 254 most abundant proteins was highly significant (r=0.47; P<0.0001), indicating that the samples can discriminate among proteins specifically enriched in different fractions. Consistent with the expectation that insoluble structures were pelleted and not subjected to analysis, we noted that tubulin and other axonemal component were no longer among the most abundant proteins as they were in the analysis described above. Major Vault Protein (MVP) was found to be among the top 10 proteins enriched in the soluble fraction from isolated membranellar bands compared to whole cells, MVP was also among the top 10 proteins that were depleted from the soluble fraction of cell bodies after membranellar band removal in (**Table S3**).

### Protein composition of the Membranellar Band

The Membranellar band is of particular interest because MB regeneration in *Stentor* is one of the best studied paradigms for regeneration of structures at the sub-cellular level. Although extensive microscopy and micromanipulation studies have been carried out to characterize MB regeneration (Tartar 1961), little is known about the process at a molecular level. Knowing the proteins that compose the MB will be an important pre-requisite for understanding the mechanisms by which they assemble into the final complex structure. In order to develop a list of MB enriched proteins, we began by clustering to identify distinct patterns of localization based on protein abundance profiles across the five fractions. As described in Methods, nbclust analysis, as well as visual inspection of a distance map, both indicate that the data best support three total clusters (**Supplemental Figure S3**). The result of Kmeans clustering with three clusters is shown in **Figure 4A.** It is evident by inspection that Cluster 1 represents proteins enriched in the MB compared to all other fractions.

**Figure 4.**
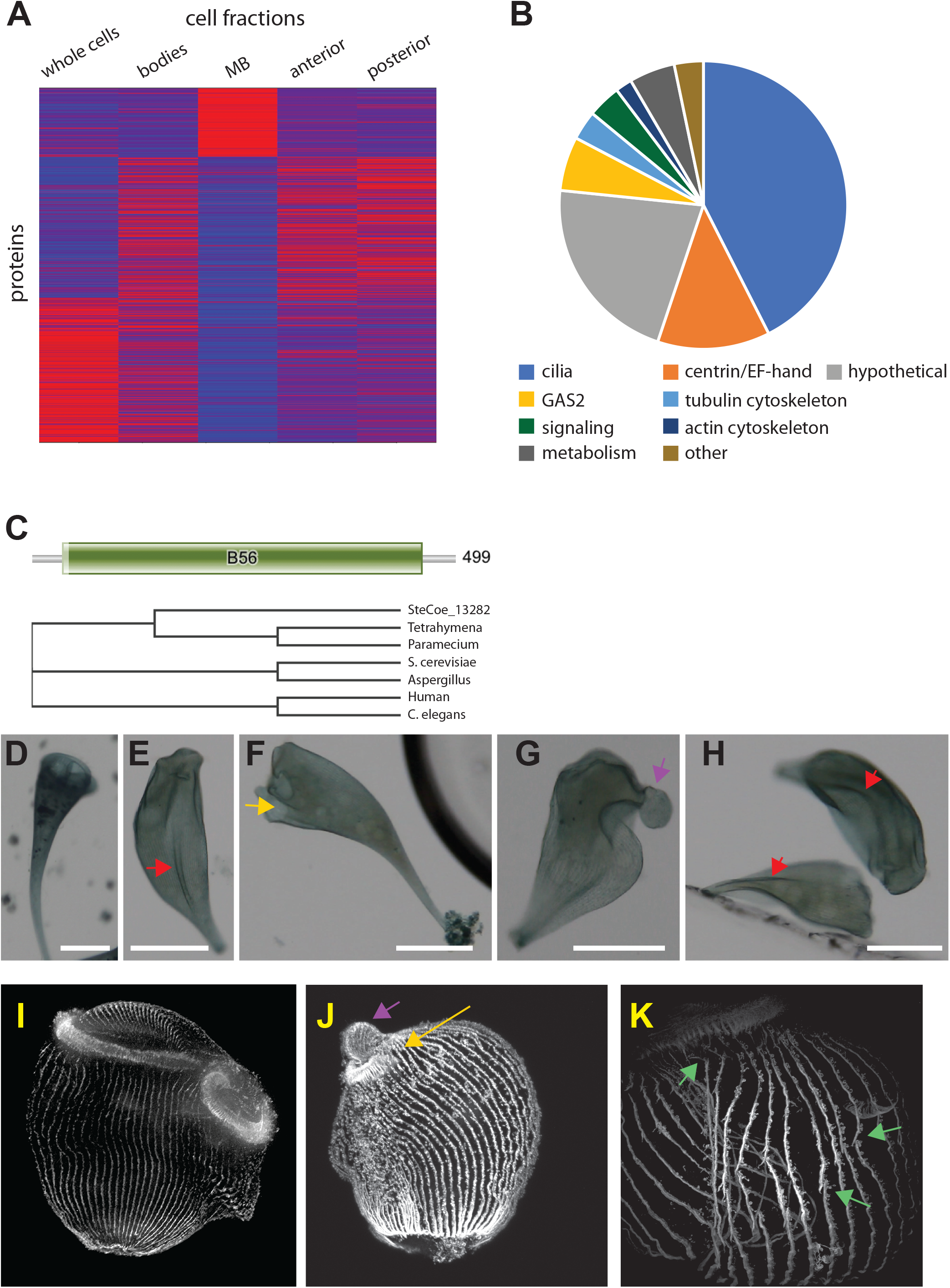
Proteome of the membranellar band. (**A**) K-means cluster analysis of abundant proteins across all five cell fragments. Cluster 1 identifies MB enriched proteins. Red indicates increased weighting in the profile. Colors indicate Z scores of normalized, median centered data for each row. (**B**) Protein families contained in the MB enriched dataset (**Table S4**) obtained by merging cluster 1 of panel A with the statistically enriched MB proteins identified in **Figure 2B** that also had RNAseq support as described in the text. **(C)** Signaling proteins of the MB include a putative PP2A B59 subunit b ortholog encoded by gene SteCoe_13282. Bar diagram shows result of PFAM domain analysis, indicating that the majority of the protein is a B59 domain characteristic of the PP2a regulatory subunit b. Phylogenetic tree is a cladogram of a Clustal multiple alignment between the stentor PP2a subunit b ortholog and PP2a subunit b from ciliate, fungal, and animal species, showing a clear grouping with the ciliate orthologs. (**D-H**) RNAi of SteCoe_13282. Scale bar 200 μm in all panels. (**D**) negative control showing the normal cone-like shape of a *Stentor* cell with a round membranellar band at the anterior end. (**E-H**) Examples of RNAi cells imaged after 9 days of RNAi by feeding. (**E**) Cell lacking a closed MB and having abnormal folds or creases on the cell surface (red arrow). (**F**) Cell in which an MB is present but where the ends do not meet up to produce a closed structure (yellow arrow). (**G**) Cell lacking a closed MB, in which the frontal field (the region of cortex normally contained inside the MB) is protruding from the cell (purple arrow). (**H**) Additional cells lacking a normal closed MB and with longitudinal folds on the cell surface (red arrows). (**I-K**) Immunofluorescence of cortical microtubule structures in PP2A RNAi. (**I**) Negative control illustrating normal cell morphology with parallel unbroken rows and a densely staining membranellar band. (**J**) RNAi cell showing a small MB with an opening on one side (yellow arrow) and a protruding frontal field (purple arrow). (**K**) Higher magnification view of cortical rows in an RNAi cell showing regions in which the cortical rows fail to maintain integrity (green arrows).

In order to develop a list of high-likelihood MB-specific proteins, we started with the proteins assigned to Cluster 1 (93 proteins) and removed two proteins that had equal abundance in MB and in cell bodies from which the MB had been removed, yielding 91 proteins. Since our clustering was based on abundant proteins, it will likely have missed enriched proteins that happened to be less abundant. It is harder to reliably calculate an enrichment value for less abundant proteins, hence we applied a set of multiple criteria to add additional proteins to our MB protein list. As detailed in methods, we applied a joint criterion in which we required proteins to be enriched in the MB sample relative to intact cells, depleted in cell bodies from which the MB was removed, and encoded by genes that were upregulated during MB regeneration based on our previous RNAseq results (Sood 2021). This additional set contained 123 proteins. Combining the two lists gave our final MB specific protein list of 214 proteins (**Table S4**). The protein classes represented in this list are plotted in **Figure 4B**.

The MB contains long, densely packed motile cilia, and it is therefore not surprising that the list contains 91 known cilia-specific proteins, including not only inner and outer dynein arm proteins but also components of the Radial Spoke complex (RSP 1, 3, 4, 7, 9, 10, 16), the Central Pair complex (Hydin, CPC1, and PF16/SPAG6), the inner junctional complex (PACRG and FAP20), the dynein regulatory complex (DRC11), and the Intraflagellar Transport machinery (IFT 56, 57, 80, 81,88, and 271).

In addition to the large group of cilia related proteins, which we had expected to see a priori, two other groups of proteins that are heavily represented in the MB specific protein list are 27 EF-hand proteins and 14 GAS2 proteins. Two of the proteins that we have annotated as GAS2-related do not contain GAS2 domains recognized by PFAM, but they show high homology to other *Stentor* genes that do contain GAS2 domains. We therefore refer to these as “GAS2 related”.

The MB specific gene list also contained eight putative proteins involved in signalling, including a CAMK-related kinase with an N-terminal ADK domain (SteCoe_29713), a CDPK-like kinase with ADK domain (SteCoe_9200), three additional CDPK family kinases (SteCoe_11641, SteCoe_20046, and SteCoe_22297), one Arrestin domain protein, one PP2C ortholog, and a Protein phosphatase 2 a (PP2a) regulatory subunit B ortholog (SteCoe_13282).

Two of these eight signaling proteins contained ADK domains in addition to their kinase domains. Tandem protein kinase ADK proteins have previously been noted as an unusual feature of the *Stentor* kinome (Reiff 2017). In addition to the two tandem ADK kinases, the MB specific proteome contains three addition proteins with ADK domains (SteCoe_15389, SteCoe_33155, and SteCoe_34629). The proteomic data also revealed two additional ADK domain containing proteins (SteCoe_15389 and SteCoe_4096) which were clearly enriched in the MB fraction but were not abundant enough to have been included in the MB specific list. ADK proteins were not similarly abundant in any other cell fractions. In total, the cobmined proteomic analysis detected eight predicted ADK domain containing proteins, all but one of which (SteCoe_7665) were enriched in the MB fraction.

### Functional analysis of MB-enriched signaling proteins reveals a PP2a subunit involved in MB morphogenesis

Among the proteins enriched in the MB, putative signaling proteins are of particular interest because they have the potential to play a role in regulating MB regeneration, either by sending signals to suppress MB formation when the normal structure is present, or by regulating steps of MB morphogenesis and maintenance. We used RNAi to target the unusual tandem ADK domain containing CAMKL and CDPK kinases enriched in the MB proteins, as well as the PP2a subunit B ortholog. Neither kinase tested showed an observable phenotype, either in vegetatively growing cells or in cells regenerating following sucrose shock. In contrast, RNAi of PP2a subunit B (**Figure 4C**) showed a dramatic morphology defect in which the MB failed to close into a ring (**Figure 4E-H**). As the phenotype develops, the frontal field often appeared to be squeezed out from the front of the cell (**Figure 4G**) and the cell surface developed deep grooves and ridges (**Figure 4E**). The same phenotypes were observed in cells treated with the PP2a inhibitor Calyculin (**Supplemental Figure S4**). Immunofluorescence imaging of cortical tubulin in PP2A RNAi cells (**Figure 4J**) shows the opening of the MB and protrusion of the frontal field, although surface folds are not seen, presumably due to contraction of the cell body that occurs during fixation. Immunofluorescence images also demonstrate that in the RNAi treated cells, the cortical microtubule rows lose their continuity, showing regions of apparent breakage (**Figure 4K**).

## Discussion

### Fraction of proteins showing polarized localization

The main result of our proteomic dissection is that approximately 30% of proteins in the *Stentor* proteome show differential localization along the anterior-posterior axis, showing enrichment in either the MB, the anterior half of the body, or the posterior half of the body. Given that our dissection into essentially three different A/P regions was relatively crude, and given that localization along other body axes was not considered, we anticipate that our results are significantly under-estimating the fraction of localized proteins.

Sub-cellular proteomics has previously been carried out by fractionating cells into distinct organelle fractions. One such study (Foster 2006) used protein correlation profiling among a series of subcellular fractions of liver cells, representing different organelles, to identify proteins uniquely localizing to different organelles by comparing the profiles of abundance of each of 1900 proteins among the fractions to reference data for nine different organelles based on known organelle specific markers. In that analysis, out of 1900 proteins, 1,258 proteins (66%) matched to at least one of the 9 organelles. The mapping of proteins to organelles was not, however, 1-to-1. Out of 968 proteins localized to one or more organelle, 373 (39%) were localized to more than one organelle according to this analysis. Similarly, fractionation profiling of lysed HeLa cells (Itzhak 2016) found that 33% of proteins had broad or cytoplasmic localization while 77% localized to one or more specific organelles. Fractionation profiling of cellular fractions from brain tissue (Pandya 2017) and from Arabidopsis cells (Dunkley 2006) indicated a similar degree of protein localization among various organellar fractions. Consistent with the high percentage of proteins localizing to distinct cellular structures in these fractionation experiments, systematic analysis of mRNA localization in Drosophila embryos revealed that more than 70% of expressed genes showed subcellular transcript localization (Lecuyer 2007). Large scale systematic immunostaining in U2OS cells was used to localized over 12,000 proteins detected by over 13,000 antibodies (Thul 2017). Image analysis was used to map each protein relative to 30 different organelles or other cellular structures. Roughly a third of the proteins in this massive study were uniformly localized in the cytosol while the other two thirds were localized to one or more distinct structures. These proteomic and cell-atlas studies thus give a consistent picture that a majority of proteins show localization to distinct subcellular structures. However, these studies were focused on co-localization with organelles, and did not address localization relative to larger scale cell polarity axes.

The fraction of proteins with polarized localization in our hands is smaller than the fraction of proteins having specific organelle localization in these prior studies. Only a sub-set of organelles appear to have polarized localizations in *Stentor* - for example, mitochondria show clear enrichment in the anterior half, but we see no proteomic signature of ER enrichment in any fraction relative to the others. Many organelles may have a radial rather than anterior-posterior localization, for example some being attached to the cortex and others more internal, and our analysis would have missed such a pattern since we only considered differences along the anterior posterior axis. Moreover, the extensive cytoplasmic streaming in *Stentor* (Slabodnick 2013) may spread many organelles throughout the cell body, such that proteins associated with those organelles will show a uniform distribution among our dissected cell pieces. Therefore, our results are not inconsistent with a high fraction of proteins localized to distinct organelles, but may simply reflect the fact that not all organelles show an anterior-posterior polarity.

The specific question of protein localization relative to cell polarity axes has been addressed using the budding yeast *S. cerevisiae.* In a systematic analyses of GFP-tagged yeast proteins (Huh et al., 2003), approximately 2% of proteins localized to the bud neck, a region of the cell surface defining the cell polarity axis in yeast. An earlier study using genome-wide epitope tagging (Kumar 2002) reported that less than 1 % of proteins localized to the bud. In a more recent analysis of GFP-tagged proteins supported by machine learning (Chong 2015) it was reported that 4% of proteins localized to either the bud, bud neck, or bud site. These three prior studies of localization in yeast used vegetatively growing cells, and looked for polarized localization relative to the bud neck. A different polarized growth process in yeast, albeit one that shares much of the same underlying polarization machinery, is formation of a mating projection in response to alpha factor. An analysis of 4200 GFP tagged clones in alpha factor treated yeast (Narayanaswamy 2009) revealed 74 proteins (1.8% of the total) with polarized localization relative to the tip. All of these systematic studies of yeast protein localization found that a number of proteins were specifically localized to anterior structures, while none were localized to the opposite pole. This draws a potentially interesting parallel to the result from our proteomic dissection of *Stentor,* in which we found that far more proteins are enriched in the anterior of the cell while relatively few are enriched in the posterior.

Another cell type that has been used to study global protein localization is the polarized epithelial cell. A comparative proteomic analysis of apical and basolateral surface proteins in Madin-Darby canine kidney (MDCK) cells identified 313 plasma membrane proteins, of which 38% were apical, 51% were basolateral, and 11% were nonpolarized (Caceres 2019). A similar study of human pluripotent stem cell-derived cyst (hPSC-cyst) cells (Wang 2021) found that out of 611 proteins identified, 59% were apical, 36% basal, and 5% nonpolarized. Thus, both of these studies, in contrast to the previous studies of protein localization in yeast, found a very large fraction of proteins were polarized.

Our results for *Stentor* thus lie between the extreme values for the fraction of the proteome that has a polarized location, ranging from a few percent in yeast cell-wide to 90% in specific comparisons of the highly polarized apical versus basolateral membranes in mammalian epithelial cells. It is interesting to consider what mechanism may allow for establishment of polarized protein localization despite cytoplasmic streaming. We speculate that because the cell cortex is covered with parallel microtubule bundles, kinesin or dynein motors may play a role in moving proteins along the cortex to one end or the other of these bundles.

### Comparison with other proteomic studies on ciliates

In *Tetrahymena,* a protein analysis of the oral apparatus (equivalent to the MB in *Stentor)* using 2D gel electrophoresis revealed 162 polypeptides (Gavin 1980) many of which had positions on the gel corresponding to components of the ciliary axoneme. Our results are consistent with that report, in that 47 out of 214 MB specific proteins are known components of axonemal structures.

A recent proteomic analysis of isolated membranellar bands from *Stentor* (Wei et al., 2020) reported the most abundant class of proteins were mitochondrial proteins, and the second most abundant class were ribosomal proteins. In our data, mitochondrial proteins are on average 3.3 fold enriched in the cell body compared to the MB (out of 53 annotated mitochondrial detected). Only 9/53 putative mitochondrial proteins were more abundant in the MB than in MB-less cell bodies, and in all cases these proteins were of low abundance. Within our MB specific protein list (**Table S4**), we find only a single protein annotated as a mitochondrial protein. Likewise, we find that ribosomal proteins are on average 2.8 fold enriched in the cell body compared to the MB (out of 141 ribosomal proteins detected). Only 12 of the 141 proteins were more abundant in the MB than in the MB-less cell bodies, and in all 12 cases these were low-abundance proteins, represented by a single peptide detected in the MB fraction. If we consider just the proteins that were detected by at least five peptides among the bodies and MB samples (47 proteins), only two were more abundant in the MB sample than in the bodies lacking MB. We conclude that both mitochondria and ribosomes are in fact depleted in the MB sample compared to the rest of the cell.

We speculate that the difference between our results and those of Wei et al (2020) may reflect methodological difference between the two studies. Wei et al. analyzed the proteome of isolated MB in order to identify abundant proteins within the sample, but did not compare the results to the proteome of the remainder of the cell. In contrast, our comparative approach allowed us to discriminate proteins based on relative enrichment between different samples. A second difference is that we removed the MB by sucrose shock, whereas Wei et al. used urea, a powerful chaotropic agent, to release MB from the cells. Given the capacity of urea to disrupt biological structures, there is the potential that the urea treatment used by Wei et al. may have led to cell lysis during MB isolation.

### Membranellar band proteins - GAS2, ADK, MVP, and PP2a

Both the membranellar band and the cortical ciliary rows are composed of motile cilia. The cilia of the MB are longer and more densely packed than those of the cell body, and this difference might in itself cause an enrichment of cilia-related proteins, relative to total protein, in the MB compared to the rest of the cell. Are there any proteins besides ciliary proteins that are specific to the MB? One notable proteomic difference that we observed between the MB and the cell body is the abundance of GAS2 orthologs. GAS2 (growth arrest specific 2) is a family of proteins with putative functions in linking actin and tubulin cytoskeletons together (N Zhang 2021). Of the 15 GAS2 related proteins (12 GAS2 domain proteins plus three GAS2 related proteins with homology to *Stentor* GAS2 domain proteins) detected in the original proteomic analysis (**Table S1**), 12 were included in the MB specific protein list, one more was detected only in the MB and not in the cell body, but was not included in the MB specific list simply because its abundance was below our cutoff. The final GAS2 in the total proteome, encoded by gene SteCoe_22878, was not detected in the MB sample. However, within the cell body samples, this protein was only detected in the anterior half and not the posterior half, consistent with it being associated with an anterior structure, possibly the MB itself. The possibility that SteCoe_22878 is a bona fide MB protein is further supported by the fact that this gene is upregulated during MB regeneration in sucrose-shocked cells (Sood 2021). In the soluble fraction of isolated MB (**Table S3**), one of the most highly enriched proteins (SteCoe_13619) is one of the GAS2-related proteins also detected in the MB specific protein list. One known function of GAS2 proteins is linking actin to microtubules, and in that light we note that three actin orthologs are found in the MB specific protein list (**Table S4**). However, the role of actin in ciliates is generally not well understood.

In the TMT comparison of anterior and posterior half-cells lacking the MB (**Figure 3**), 12 GAS2 proteins were identified as being differentially enriched between the anterior and posterior halves. Out of these 12, 11 were enriched in the anterior half. Four of these 11 proteins were also contained in the MB specific protein list. Because the TMT analysis was done on bisected cell bodies from which the MB had been removed, these proteins are evidently not strictly specific for the MB, even though they are highly enriched in that sample. This may reflect incomplete removal of the MB from the cell bodies during sucrose shock, or it may indicate a true dual localization of these particular GAS2 proteins in the MB and in some other anterior structure.

Another family of proteins highly represented in the MB specific protein list are the adenylate kinase (ADK) domain containing proteins. Seven of the eight ADK domain proteins identified in our proteome were enriched in the MB. ADK is involved in ATP regeneration in motile cilia in some organisms (Kinukawa 2007), so the enrichment of ADK in the MB could simply reflect the high density of motile cilia. However, the fact that ADK domains appear to be present on other MB-associated kinases raises the possibility that the ADK domain may play some role in the assembly or targeting of proteins into the MB.

The presence of major vault protein (MVP) in the MB was unexpected. MVP is a component of vaults, large ribonucleoprotein complexes that are present in many eukaryotes including humans (Rome 1991; Berger 2009). Although vaults are highly conserved, their function remains unclear. Vaults have been hypothesized to play roles in immunity, drug resistance, signaling and nucleo-cytoplasmic transport. The Vault complex consists of three protein subunits. MVP forms a large 670 angstrom barrel (Tanaka 2009). Inside the barrel are two accessory proteins, vault poly(ADP)ribo polymerase (vPARP; Kickhoefer 1999a) and telomere associated protein (TEP; Kickhoefer 1999b). Our proteomic analysis detected two PARP orthologs neither of which was enriched in the MB fraction, and did not detect any TEP orthologs. One possibility is that the MVP orthologs in *Stentor* assemble into a structure unique to the MB, rather than canonical vault structures seen in other species. RNAi experiments with *Stentor* MVP genes did not reveal any observable phenotypes.

The RNAi analysis of **Figure 4D-K** indicates that the MB-enriched PP2a plays a role in proper MB morphogenesis. We did not observe any evidenced of ectopic MB formation as would be expected if the protein was part of a pathway that inhibits regeneration. Instead, the defects appear to be morphological. PP2a interacts genetically with the Mob1 client kinase NDR1 in fungi (Shomin-Levi 2017). Given our previous result that RNAi of Mob1 in *Stentor* results in cells with multiple membranellar bands (Slabodnick 2014), the new result with PP2a further points to the Mob1/NDR pathway as playing an important role in MB morphogenesis.

### Implications of temporal expression pattern of MB-enriched genes

We previously reported an RNAseq analysis of gene expression during MB regeneration in Stentor (Sood 2021). In that study we defined two lists of genes upregulated during MB regeneration. One list compared regeneration following removal of the MB by sucrose shock with regeneration in the posterior halves of bisected cells, which have to regenerate an MB, and identified genes upregulated in both cases, resulting in what we term the OA (oral apparatus) specific gene list. A second list consists of genes upregulated during sucrose shock regeneration but not in regeneration of bisected cells, which we refer to as the sucrose specific gene list. Out of the 214 proteins in the MB specific protein list (**Table S4**), 168 are upregulated during regeneration of the MB (28 in the OA gene list, 132 in the sucrose shock gene list, and 5 in the anterior regeneration gene list). Each of these lists were then clustered into five temporally distinct waves of gene expression. The largest number of genes encoding MB specific proteins in our dataset fell into expression Cluster 4-5 of the OA specific gene list (defined by having peak expression at 5-6 hours) and clusters 3-4 of the sucrose shock gene list (peak expression at 3-4 hours). Sucrose shock clusters 3 and 4 were previously noted to contain a large number of cilia-specific genes, consistent with the abundance of cilia related proteins in the MB specific protein list. The timing of expression of the MB specific proteins is substantially later than the time period during which the cilia first assemble, but instead corresponds to the time at which MB cilia begin coordinated motility to generate long-range fluid flows (Paulin and Bussey 1971; Wan 2020). These results are consistent with our previous observation that genes whose products are involved in ciliary motility show a generally later expression in *Stentor* regeneration than genes involved in building the cilium itself (Sood 2021).

The MB-specific GAS2 proteins are expressed in clusters OA4, O5, and Sucrose 4, all of which peak long after the time of basal body assembly but coincide with the time that basal bodies, which originally form in a random “anarchic field” (Bernard 1981) become organized into membranelles, and the cilia begin to acquire coordinated beating.

In contrast to the large fraction of MB specific proteins whose genes are upregulated during regeneration, only 4/206 differentially localized anterior or posterior proteins (based on TMT analysis **Table S2**) are upregulated during regeneration of bisected *Stentor coeruleus* cells. Both of the posterior-specific centrins (SteCoe_ 9428and SteCoe_3266) show maximal expression 7 hours after bisection (Sood 2021). This is long after the completion of holdfast regeneration, which takes place within the first two hours of regeneration (Morgan 1901; Weisz 1951). 26 of the anterior/posterior differentially localized proteins in **Table S2** correspond to genes upregulated in the RNAseq analysis of regeneration in bisected cells of the related ciliate *Stentor polymorphus* (Onsbring 2018), of which the majority (22 out of 26) show peak expression 5 hours after bisection, by which time the bisected cell halves have restored an overtly normal morphology. Thus, judging by transcriptional timing in both species, it appears that proteins with polarized localizations in the cell body show peak synthesis long after cell morphology has been re-established, implying either that these proteins are not required for morphogenesis, or that sufficient reserves of these proteins were left in the cell fragments to support proper regeneration.

### Proteomic Dissection as a strategy for Spatial Proteomics

These results show that it is possible to map intracellular protein distributions by physically cutting a cell into pieces and then analyzing the fragments. The unusually large size of *Stentor* enables this strategy to be implemented using manual dissection with a glass needle, but such an approach will not work with smaller cells. However, a variety of microfabricated cutting devices capable of working at the cellular level are now being developed (Blauch 2017; Koppaka 2021; K Zhang 2021). This growing ability to cut tiny pieces of cells and tissues, combined with increasing sensitivity for proteomic analysis of small samples such as a single mammalian cell (Dou 2019; Zhu 2019; Tsai 2020; Schoof 2021), means that proteomic dissection, as demonstrated in this work, can provide a general method for subcellular proteomics that is complementary to existing methods that based on fractionation or imaging.

## Acknowledgments

We thank the members of the Marshall lab, as well as Susan Fisher and Joana Caldeira for helpful discussion and suggestions. This work was funded by NIH grant R35 GM130327 (WFM), by an NSF graduate research fellowship (AL), and by the HHMI Gilliam Fellowship (UD). WFM is a Chan Zuckerberg Biohub investigator. Portions of the research were supported by grant U24CA210955 from the National Cancer Institute and grant P41 GM103493 from the National Institute of General Medical Sciences. Some of the work was performed in the Environmental Molecular Sciences Laboratory (grid.436923.9), a U.S. Department of Energy (DOE) Science User Facility at the Pacific Northwest National Laboratory (PNNL) sponsored by the Office of Biological and Environmental Research. PNNL is a multiprogram national laboratory operated by Battelle for the DOE under contract DE-AC05-76RL01830

## Author Contributions

AL dissection experiments, data analysis, made figures

PDP mass spectrometry, data analysis

CFT mass spectrometry, data analysis

LY mass spectrometry, data analysis

TM RNAi experiments

UD RNAseq data analysis

CY RNai experiments

DS illustrations

PS RNAseq data analysis

RDS mass spectrometry methods

TL mass spectrometry, data analysis, wrote paper

WFM designed experiments, data analysis, manual annotation, made figures, wrote the paper

---

**Supplemental Figure S1.**
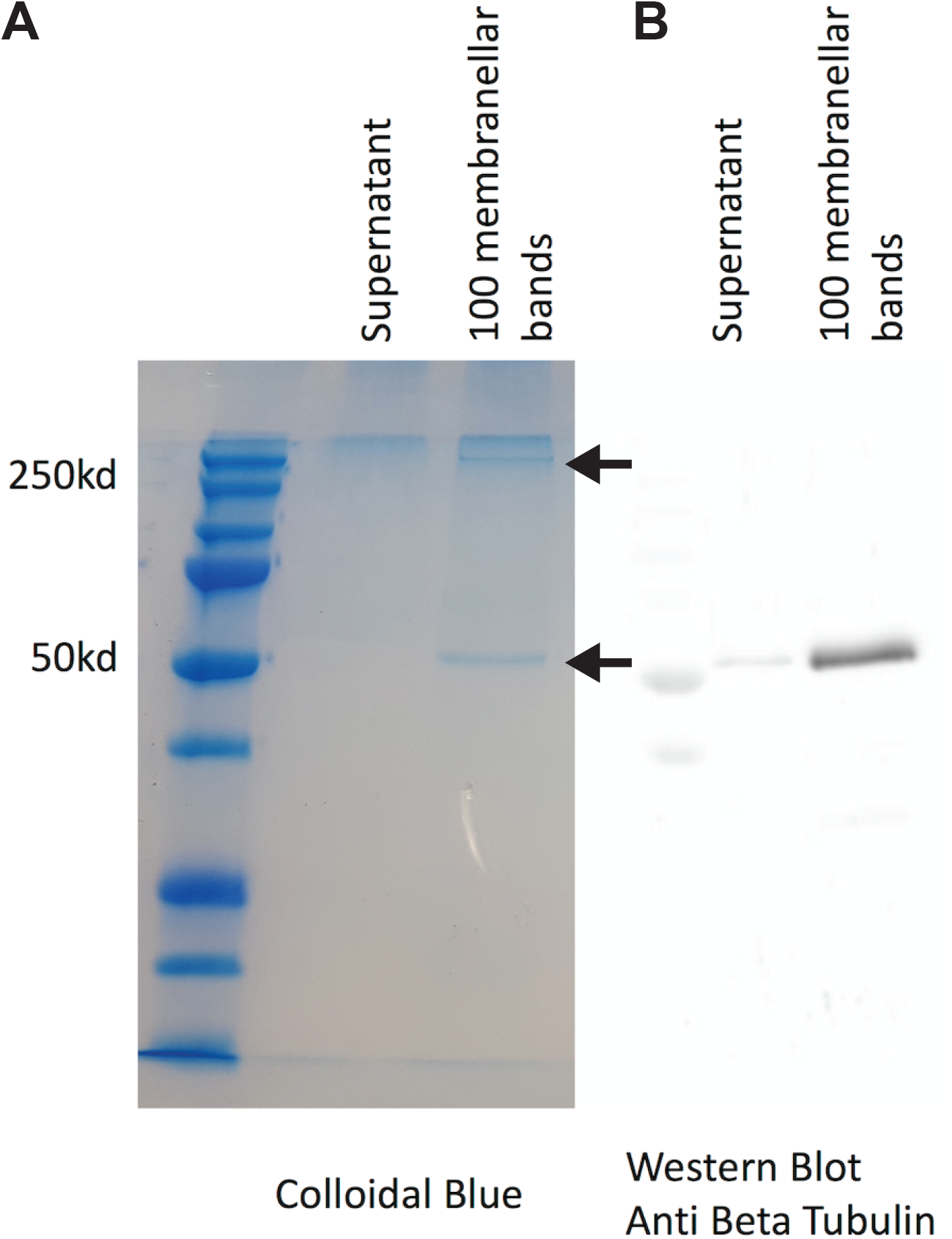
Identifying most abundant proteins in MB fraction. (A) Colloidal blue staining of isolated MB along with supernatant obtained during the MB collection. (B) Western blot for tubulin confirming the identity of the 55 kDa band.

**Supplemental Figure S2.**
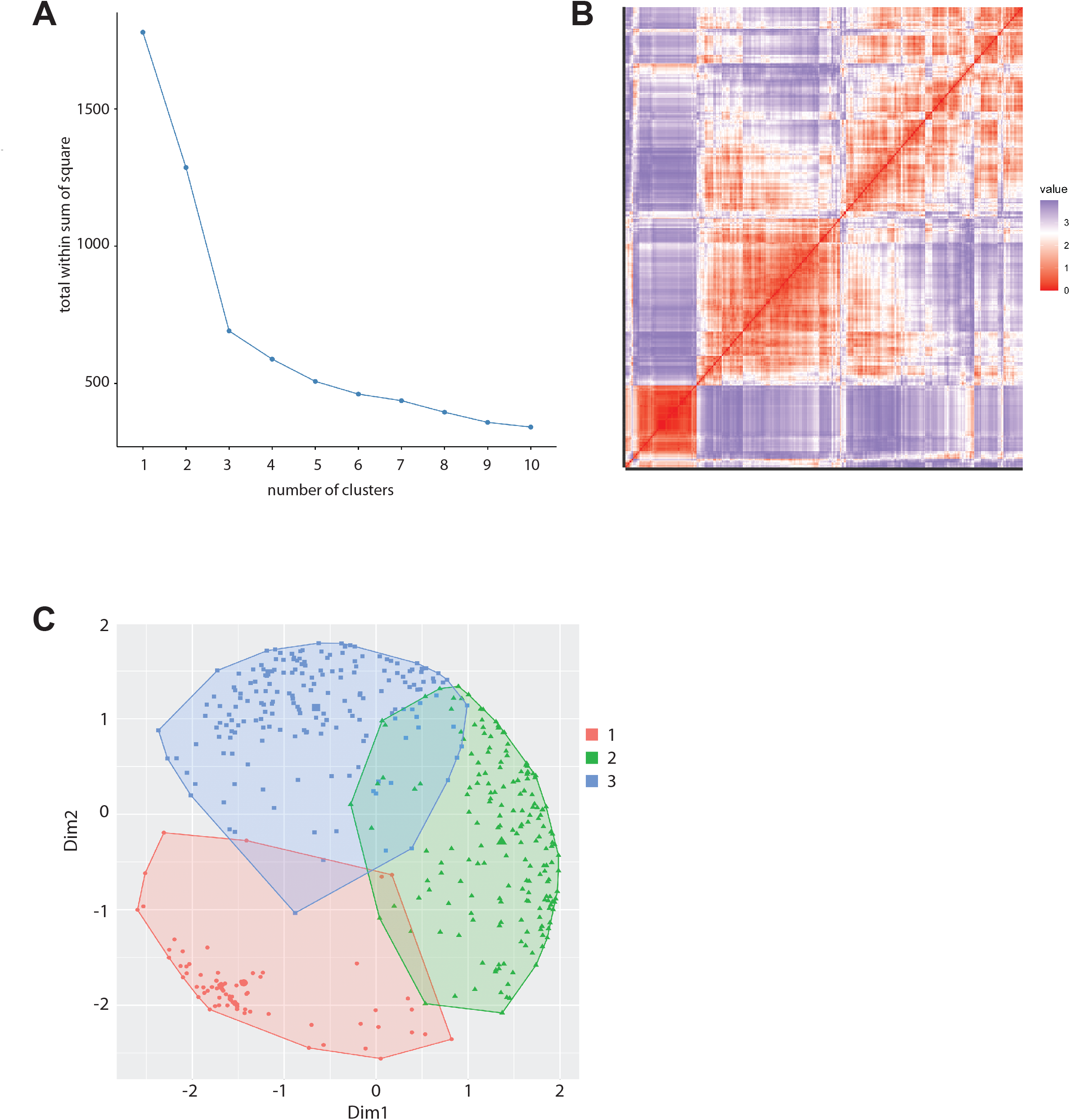
Pairwise comparison between samples. Each pie chart represents the five proteins most enriched in each *Stentor* sample compared to each other sample, regardless of total quantity. Color scheme is the same as in **Figure 2C**.

**Supplemental Figure S3.**
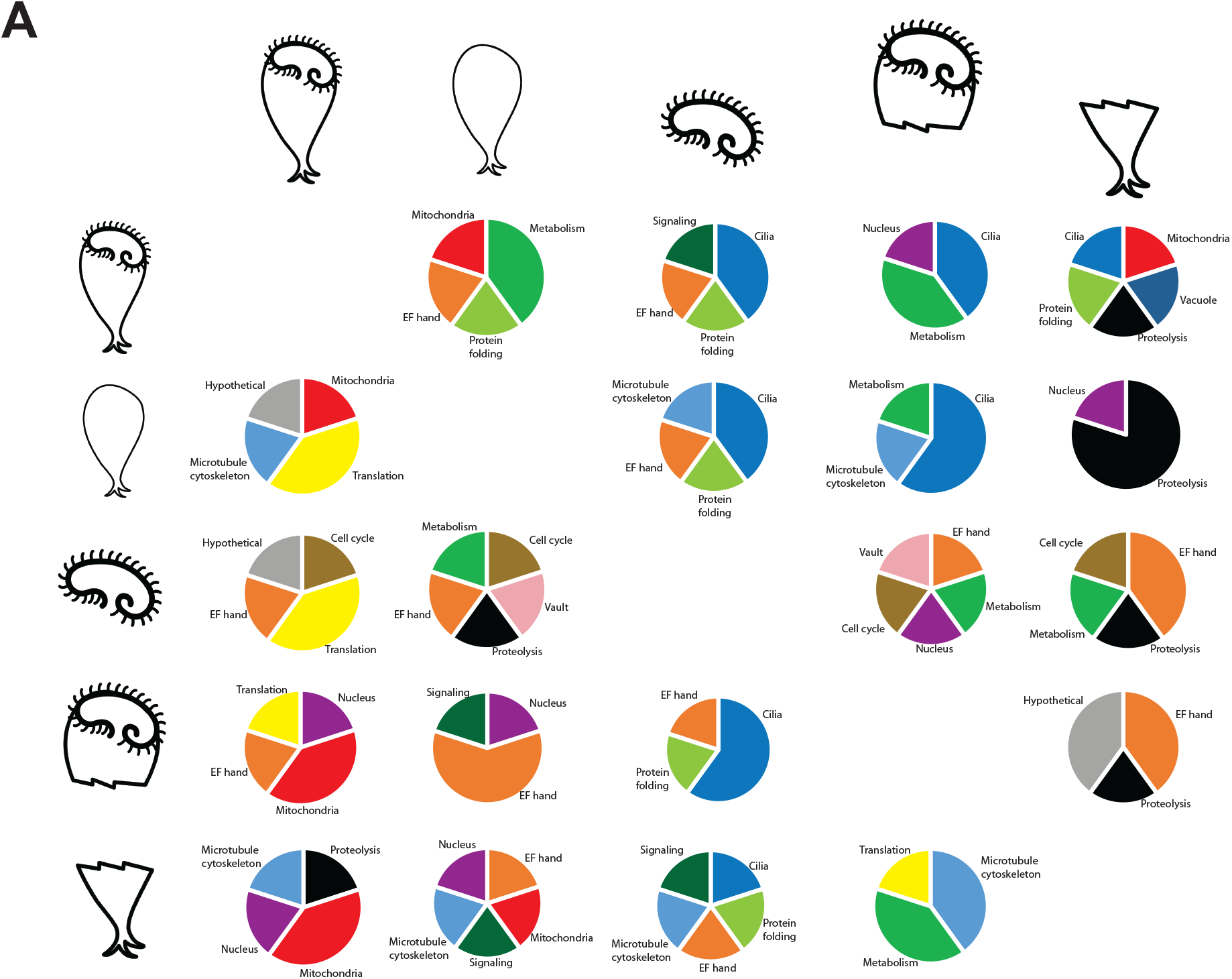
Clustering of proteomic data. (A) Nbclust plot showing clear inflection point at 3 clusters. (B) Distance map showing three blocks of correlation. (C) Cluster diagram from same assignment as used to produce the heatmap in **Figure 4A**, indicating that cluster 1 is well separated from clusters 2 and 3.

**Supplemental Figure S4.**
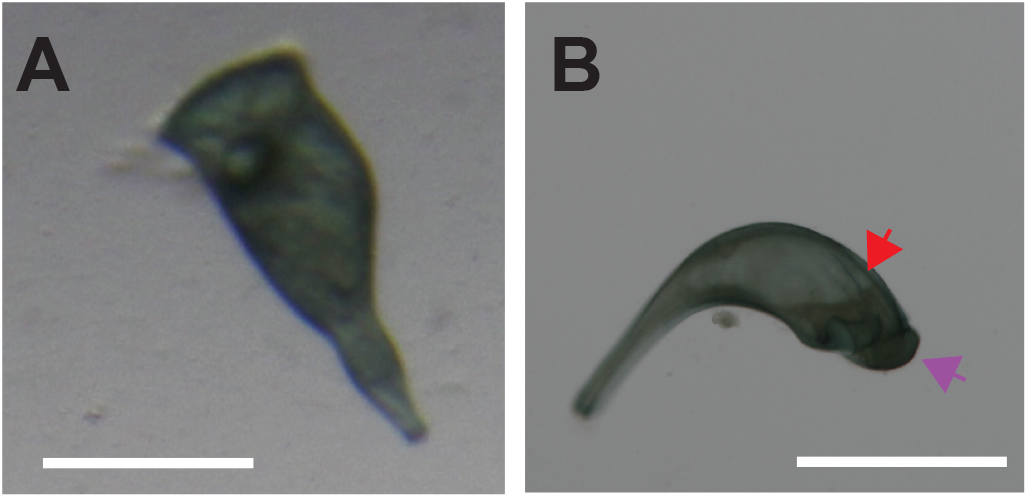
Calyculin A treatment recapitulates phenotype of PP2a RNAi. (**A**) Untreated cell showing normal morphology. (**B**) Calyculin A streated cell showing surface folds (red arrow) and protruding frontal field (purple arrow), comparable to phenotypes seen with PP2a RNAi in **Figure 4**.

## Methods

### Cell culture

*Stentor* from Carolina Biological Supply Company (Burlington, NC) were grown at room temperature in pasteurized spring water and fed *Chlamydomonas,* wild type CC125, every 5 days.

### Preparation of Stentor membranellar bands and cell fragments

*Stentor* sucrose shocking and surgery were performed as previously described (Lin 2018). Briefly, whole *Stentor* were collected in 5ul of modified *Stentor* media (MSM) consisting of 0.75 mM Na_2_CO_3_, 0.15 mM KHCO_3_, 0.15 mM NaNO_3_, 0.15 mM KH_2_PO_4_, 0.15 mM MgSO_4_, 0.5 mM CaCl_2_, 1.47 mM NaCl (Slabodnick 2017). *Stentor* were slowed by placing them in 2% methylcellulose and dissected using glass needles pulled from capillary tubes. Methylcellulose was subsequently washed out and *Stentor* halves were collected in 5ul of MSM. *Stentor* were sucrose shocked in 12% sucrose, membranellar bands were spun down and supernatant was removed. Bodies were washed and collected in 5ul MSM.

### Gel analysis of isolated MB

For gel electrophoresis, MB were collected by urea shock with 4% urea in MSM. Final samples were dissolved in 1% SDS, 0.05M Tris, 0.12mg w/v bromophenol blue, 6% glycerol and 5% betamercaptoethanol. before running on a 10% SDS gel at 200V, 0.1A, 50W, for 45 minutes. For proteomic analysis of bands, gels were stained with colloidal blue (**Supplemental Figure S1A**). Cut bands were frozen and analyzed by mass spectrometry (Applied Biomics Inc). In the 250 kDa band, there were 18 proteins identified, mostly with just one or two peptides. In contrast, a protein corresponding to IDA1 inner dynein arm heavy chain (SteCoe_11197) was identified with 17 peptides, supporting the identity of the 250 kDa band as a dynein heavy chain protein. To confirm tubulin band identity, Western blotting was performed using overnight transfer at 4 degrees C, at 16mA onto Immuno-Blot PVDF Membrane (Bio-Rad). Blots were blocked 1 hr., stained with monoclonal anti-alpha tubulin mouse antibody (1:1000 dilution) for 1 hour, washed three times for 10 min in PBS, stained 1 hour AlexaFluor 546 goat-anti mouse IgG (1:1000 in PBS), and washed three times for 10 min in PBS. (**Supplemental Figure S1B**).

### Mass Spectrometry

The *Stentor* cell samples (intact cells or membranellar bands) were prepared as described previously (Huang et al., 2016), with proteins being solubilized using 8 M urea for analysis of total protein or 0.1% n-Dodecyl β-D-maltoside (DDM) for analysis of soluble proteins, in 50 mM TRIS pH 8. For each sample, approximately 25 cells worth of material was loaded for analysis. The resulting protein extracts were analyzed using the simplified nanoproteomics platform (SNaPP), a custom online digestion system described in previous publications (Piehowski et al. 2018, Clair et al., 2016). Briefly, the sample was passed through a 150 μm I.D. fused silica capillary packed with Poroszyme (Thermo Fisher) immobilized trypsin, at 0.5 μL/min to achieve digestion. The resulting peptides were desalted inline by trapping on a 150 μm ID × 4 cm C18 column. After washing, peptides were separated on an in-house packed 50 μm ID × 75 cm C18 analytical column. The SNaPP system was coupled to a QExactive Plus mass spectrometer (Thermo Scientific) operated in a top-12 data-dependent acquisition mode, with a 100-ms MS2 maximum ion injection time to increase sensitivity. LC-MS/MS data was searched using MSGF+ (Kim and Pevzner 2014) against the *Stentor coeruleus* protein sequence database (November 2016 release downloaded from http://stentor.ciliate.org/). Decoy database search was used to filter datasets to a 1% false discovery rate at the unique peptide level. Confident peptide-to-spectrum matches (PSMs) were summed to create protein-level spectral counting results.

For proteomic comparison of proteins between top and bottom halves of MB-less *Stentor* cell bodies, protein from top and bottom of *Stentor* cell bodies were extracted by 0.2% DDM in 50 mM HEPES at 80 °C for 50 mins. The sample solutions were diluted to 0.02% DDM digested with 1 μg lysyl endopeptidase (Wako) for 3 h followed by 1 μg trypsin (Pierce) at 37 °C overnight. After digestion, the peptides were desalted by reversed phase-Stage Tips (Rappsilber, 2007) and the concentration of peptides were estimated via BCA assays. Then, the tryptic peptides from top and bottom of Stentor cell were labeled isobarically with the 6-plexed tandem mass tag (TMT) reagents (ThermoFisher Scientific): TMT126 (top), TMT127 (bottom), TMT128 (top) and TMT129 (bottom), respectively, as previously described (Tsai et al., 2019) (The TMT to peptide amount was around 25:1). The TMT-labeled peptides were acidified to stop the labeling reaction and diluted to reach a final acetonitrile concentration of 4%, after which they were desalted by reversed phase-Stage Tips (Rappsilber, 2007) and fractionated and concatenated into 12 fractions using nanoFAC (nanoflow Fractionation and Automated Concatenation) (Dou et al., 2019). The final concatenated fractions were analyzed by QExactive Plus mass spectrometer as previously described (Dou et al., 2019). Briefly, data were acquired in a data-dependent mode with a full MS scan from m/z 350-1800 at a resolution of 70,000 at m/z 400 with automatic gain control (AGC) setting set to 3×10^6^ and maximum ion injection period set to 100 ms. Top-10 precursor ions having intensities >5×10^3^ were selected with an isolation window of 1 Da for MS/MS sequencing at a higher-energy collisional dissociation (HCD) energy of 35%. The MS/MS spectra were acquired at a resolution of 17,500. The AGC target was 2×10^5^ and the maximum ion accumulation time was 300 ms. The dynamic exclusion time was set at 40 s. LC-MS/MS data was searched using MSGF+ and the TMT reporter ion intensities were extracted using MASIC (Monroe et al., 2008). The TMT intensities from peptides with q value <0.01 were summed to create protein-level TMT intensities results and analyzed by Perseus (Tyanova et al., 2016) for statistical analyses.

### Protein Annotation

Hits were identified using the protein predictions from gene models of *Stentor* genome v1.0 (available online at stentor.ciliate.org) as reported in Slabodnick (2017). Only proteins with at least two hits based on raw counts were retained for the analysis in **Table S1**. Relative abundances using median-centered data were used to calculate ratios between parts.

Many cilia-related proteins are known to contain functional domains such as WD40, armadillo, and leucine-rich repeats. Any proteins annotated in the *Stentor* genome as “hypothetical proteins” but containing domains common among ciliary proteins were blasted against the *Chlamydomonas reinhardtii* genome (version 5.6; Phytozome.jgi.doe.gov), in which the annotation of cilia related has been most extensively carried out based on proteomic and genetic analyses.

Because SFI motif proteins often share little homology outside of the SFI1 motifs, grep was used to detect proteins matching the regular expressions ‘LL........[FL]..W[KR]’ and ‘LL......[FL]..W[KR]’ withing all proteins identified in the proteomic analysis. These were then annotated as SFI proteins.

### Dissection Profiling

The dissection profile (**Figure 1F and G**) was calculated by comparing two samples (MB versus bodies lacking MB) to a third sample representing whole cells, using only proteins with at least 5 hits across all samples. A proportionality factor p is iterated over the range 0 to 1 in 100 increments. At each value, a weighted abundance is calculated for each protein by taking the sum of A*p and B*(1-p) where A and B are the abundances in the first two samples (MB and cell bodies). This calculation is performed across all proteins and the resulting set of weighted abundances is compared to the abundances of the same proteins in the whole cell sample using a Chi squared test (via the R chisq.test function). The optimal value of p is that which minimizes Chi squared. **Figure 1F,G** plots the value of the Chi squared statistic as a function of the proportionality factor.

### Binomial test for enrichment

**Figure 2A** plots the correlation coefficient for median weighted protein hits normalized by whole cell data, between membranellar bands, cell bodies missing the MB, anterior halves of bisected cells, and posterior halves of bisected cells.

To identify the number of proteins significantly enriched in MB versus bodies-bands (**Figure 2B**), we considered only proteins for which the sum of hits in the MB and bodies fractions was at least 5. We used the binomial distribution to calculate the probability that a given protein was significantly enriched in one or the other of the two fractions, using a cutoff probability of 0.05. To compare proteins localized in the anterior versus posterior half cells, we first subtracted all proteins that showed an enrichment (significant or not) in MB versus bodies from the total list of proteins, and then of the remaining proteins we considered only those for which the sum of raw counts in the anterior and posterior half-cell fractions was at least 5. Again, we applied the binomial test with a cutoff probability of 0.05.

### K-means Cluster Analysis

After filtering only those proteins detected at least 10 times across samples, median-centered abundances were Z-normalized and then used for clustering in R. The number of clusters was first assessed by finding a clear inflection point in a nbclust plot using the R function fviz_nbclust (**Supplemental Figure S3A**) which revealed evidence for three clusters. This was also confirmed by visual inspection of a distance map (generated using fviz_cluster) which showed three blocks of correlations (**Supplemental Figure S3B**). Assignments based on K-means clustering using the kmeans function of the R cluster package with 25 starts are plotted in a cluster diagram (**Supplemental Figure S3C**) and yielded the heatmap shown in **Figure 4A**.

### Criteria for MB specific protein list

In order to compile a curated list of proteins likely to be true components of the MB, we merged two sets of proteins. In the first set, we took the 93 proteins of Cluster 1 (see **Figure 4A**), and excluded two proteins that, by inspection, were seen to be equally abundant in the MB sample as in the headless cell body sample. This produced a revised cluster 1 set of 91 proteins. To these 91 proteins, we added a second set of proteins obtained by combining the following criteria: First, we required the proteins not to be components of clusters 2 or 3. Second, we required proteins to show at least a 10-fold enrichment in MB versus whole cell samples, while also being at least 2-fold depleted in cell bodies lacking MB compared to whole cells. Third, we employed transcriptomic support by requiring that the genes encoding the proteins are upregulated in cells regenerating the MB (clusters 2-5 of the OA, sucrose, or anterior gene lists from Sood 2021). Finally, we excluded any proteins that were enriched in the posterior half-cell compared to the anterior half-cell. Applying all of these criteria to proteins not in cluster 1, resulted in a list of 123 proteins. When this was merged with the modified cluster 1 list, a list of 214 proteins was obtained that we expect to represent bona fide MB proteins. We note that the criteria were highly selective, and this list is likely missing some proteins that are actually components of the MB, but for which the supporting evidence is not as strong.

### RNAi analysis

Primer sequences for the construct targeting PP2a subunit b were 5’ TACAGCAGGCCGAGGTAAAG 3’ and 5’ TGAGTTACCAAAAGGCCAATATC 3’. RNAi by feeding and tmmunofluorescence were performed as previously described (Slabodnick 2014). Live cells were imaged using a Zeiss Axiozoom at 40x or 80x magnification.

### Comparison of proteomic data to RNAseq data

RNAseq data for regeneration in *S. coeruleus* were taken from the supplemental tables of Sood et all 2021, and merged with the proteomic data using a left join operation on the gene accession numbers. For comparison with RNAseq information in *S. polymorphus* (Onsbring 2018), the published *S. polymorphus* data was used to assemble a transcriptome. Removal of adapter content, low quality reads, and contamination were accomplished using (FASTQC, trimomatic, CutAdapt, Bowtie2). After quality control, the reads are used to generate individual count tables for each sample(Kallisto). These count tables are then used to generate normalized read counts and statistics (DESeq2). Genes were identified that showed significant differential gene expression during regeneration of posterior half-cells of *S. polymorphus,* and clustered based on the timing of peak expression. To map the transcriptomic data from *S. polymorphus* onto the corresponding genes in *S. coeruleus,* reciprocal best hits were obtained as previously described (Moreno-Hagelsieb 2008). Results of this analysis are reported in Table S1.

